# Preserved CETP Activity and Early Serum Amyloid A are Associated with Improved Outcomes in Sepsis-Associated ARDS

**DOI:** 10.64898/2026.09.14.751535

**Authors:** Ailing Ji, Osemwenoyenmwen Abu, Luke W. Meredith, Hui Yu, Clairity H. Voy, Maria E.C. Bruno, Scott M. Gordon, Gregory S. Hawk, Marlene E. Starr, Preetha Shridas

## Abstract

**Background:** Sepsis is a life-threatening condition characterized by organ dysfunction resulting from a dysregulated host response to infection. It is a leading cause of in-hospital mortality. The multicenter Acute Respiratory Distress Syndrome Network Statins for Acutely Injured Lungs from Sepsis (ARDSNet-SAILS) trial found that rosuvastatin (RS) did not improve clinical outcomes in patients with sepsis-associated ARDS and may increase hepatic and renal dysfunction. We evaluated associations of plasma cholesterol ester transfer protein (CETP) activity and Serum Amyloid A (SAA) levels with survival and clinical outcomes using plasma samples and clinical data from ARDSNet-SAILS trial.

**Methods:** This secondary analysis included clinical data and plasma samples from 471 ARDSNet-SAILS participants, randomized to placebo (PL) or RS. Plasma CETP activities and SAA levels were measured at enrollment (Day 0, n=466) and Days 3 (n=425) and 6 (n=342). Treatment effects were assessed using linear mixed models, and associations with survival and outcomes using Cox, linear, beta-binomial, and zero-inflated negative binomial regression models.

**Results:** CETP activities were significantly higher in the survivors compared to non-survivors on days 0, 3, and 6 of analysis, in both the treatment groups (p<0.01). Higher Day 0 and Day 3 CETP activities in PL patients independently predicted improved survival, with every 3-unit increase in activity associated with 18–19% lower mortality risk. In the RS treatment group, higher CETP at Day 0, Day 3, and Day 6 was significantly associated with reduced hazard of mortality (p=0.039, 0.005, and 0.022, and hazard ratios = 0.87, 0.82, and 0.80, respectively, for Days 0, 3, and 6). Higher CETP values on Days 3 and 6 were also significantly negatively associated with APACHE score and positively associated with increased renal dysfunction-free days (p=0.001). Plasma SAA levels decreased significantly over time in both treatment groups (p<0.0001). In the PL group, higher plasma SAA levels were associated with a lower hazard of mortality. Accordingly, survivors had significantly higher plasma SAA levels than non-survivors at Day 0 (p=0.005). Higher SAA levels at Days 0 and 3 were also inversely correlated with APACHE scores, while Day 0 SAA levels were positively associated with overall organ dysfunction-free days (p=0.002). In contrast, in the RS group, higher plasma SAA levels were associated with an increased hazard of mortality, reaching statistical significance at Day 6 (p<0.01). Despite this, higher SAA levels in the RS group were positively associated with renal dysfunction-free days (Day 3, p=0.004; Day 6, p=0.006), hepatic dysfunction-free days (Days 0, 3, and 6; all p≤0.001), and overall organ dysfunction-free days (Day 0, p=0.017; Days 3 and 6, both p<0.001). Analysis of SAA levels in FPLC fractions demonstrated higher SAA levels in the HDL fraction among survivors compared with non-survivors in the PL group on Days 0 and 3; however, this difference was not observed in the RS group.

**Conclusion:** Taken together, these findings identify plasma CETP activities and early SAA levels as stable prognostic biomarkers associated with improved survival and reduced disease severity in sepsis. However, while both CETP activity and SAA levels were unaffected by RS treatment, it appeared to modify the association between SAA and mortality.

## Introduction

The Statins for Acutely Injured Lungs from Sepsis (SAILS) trial was a multicenter, randomized, PL-controlled clinical trial conducted by the ARDS Network to evaluate whether RS could improve outcomes in patients with sepsis-associated acute respiratory distress syndrome (ARDS). The rationale for statin therapy in this setting was based on evidence that statins exert biological effects beyond cholesterol reduction, including attenuation of inflammation, oxidative stress, endothelial dysfunction, and immune activation (1–3). These pleiotropic effects, together with observational studies suggesting potential benefits of statins in sepsis and ARDS, provided the foundation for evaluating RS as a potential therapy for critically ill patients.

Patients enrolled in SAILS were randomized to receive PL or RS (40 mg loading dose followed by 20 mg daily) until the third day after intensive care unit discharge, study day 28, hospital discharge, or death. Detailed descriptions of the study design, eligibility criteria, and clinical outcomes have been reported previously (4). Despite a strong mechanistic rationale, RS did not improve mortality, ventilator-free days, or other clinical outcomes in patients with sepsis-associated ARDS. Moreover, RS treatment was associated with fewer hepatic and renal failure-free days, raising concerns regarding the safety of statin therapy in the setting of severe sepsis and critical illness.

RS belongs to the statin family of drugs, prescribed for treating hypercholesterolemia. Beyond lipid lowering, statins influence multiple pathways involved in inflammation, vascular homeostasis, and innate immune regulation (3). Understanding how RS influences lipid-associated pathways during sepsis may provide insight into mechanisms underlying treatment response and identify biomarkers associated with recovery or adverse outcomes. CETP is a key regulator of lipoprotein metabolism. CETP facilitates the transfer of cholesteryl esters from high-density lipoprotein (HDL to apolipoprotein B–containing lipoproteins in exchange for triglycerides, resulting in HDL particles that are depleted of cholesterol and more rapidly cleared from the circulation (5, 6). Meta-analyses of large population cohorts, investigating the effects of polymorphisms associated with low CETP activity, have shown higher HDL-C, low low-density cholesterol (LDL-C) levels, and a reduced risk of cardiovascular disease (7, 8). The role of CETP in sepsis outcomes remains controversial. In humans, a gain-of-function CETP genetic variant has been associated with lower HDL levels, increased organ dysfunction, and higher mortality during sepsis (9). In contrast, preclinical studies in mice expressing human CETP have demonstrated protective effects against sepsis mortality (10, 11). These conflicting findings highlight the need for further investigation into the relationship between CETP, lipoprotein metabolism, and clinical outcomes in sepsis.

SAA is an acute-phase apolipoprotein that rapidly associates with HDL during inflammatory states and substantially alters HDL composition and function (12, 13). SAA has been proposed as a sensitive biomarker of inflammation in bacterial and viral infections, comparable to, or potentially exceeding, the sensitivity of C-reactive protein (CRP) (12–14). Beyond its role as a biomarker, however, the relationship between SAA levels and clinical outcomes in acute inflammatory syndromes such as sepsis remains poorly defined. Our studies have demonstrated that SAA deficiency exacerbates lung injury and increases mortality in murine models of sepsis (15, 16), supporting a potential biological role for SAA during severe systemic inflammation.

In the present study, we investigated the association between lipid-related biomarkers and clinical outcomes in patients enrolled in the SAILS trial. Using archived plasma samples collected at enrollment and on study days 3 and 6, we quantified CETP activities and SAA levels. We then evaluated the relationships between these biomarkers, RS exposure, and sepsis outcomes, including organ dysfunction and mortality.

## Methods

### Population

This is a secondary analysis of randomized clinical trial outcome data from the ARDS Network SAILS trial (4). Details of the treatment modalities and baseline characteristics of all patients, including those who received RS, are provided in the original trial publication (4). The original trial was conducted from March 18, 2010, to September 30, 2013, and enrolled 745 participants. Eligible patients had a known or suspected infection, met the systemic inflammatory response syndrome (SIRS) criteria, and had acute lung injury. Participants were randomized to receive either RS or PL. EDTA-anticoagulated blood samples were collected on Day 0 (enrollment), Day 3, and Day 6, centrifuged, and the plasma was stored at −70°C. Plasma samples and clinical data were obtained from the NHLBI Biologic Specimen and Data Repository Information Coordinating Center (BioLINCC) for use in the current study. Plasma samples received from BioLINCC on dry ice were stored at −80°C until use. Plasma samples from 471 patients were ultimately received (Fig. 1), although not all patients had samples available at all three time points.

**Fig. 1.**
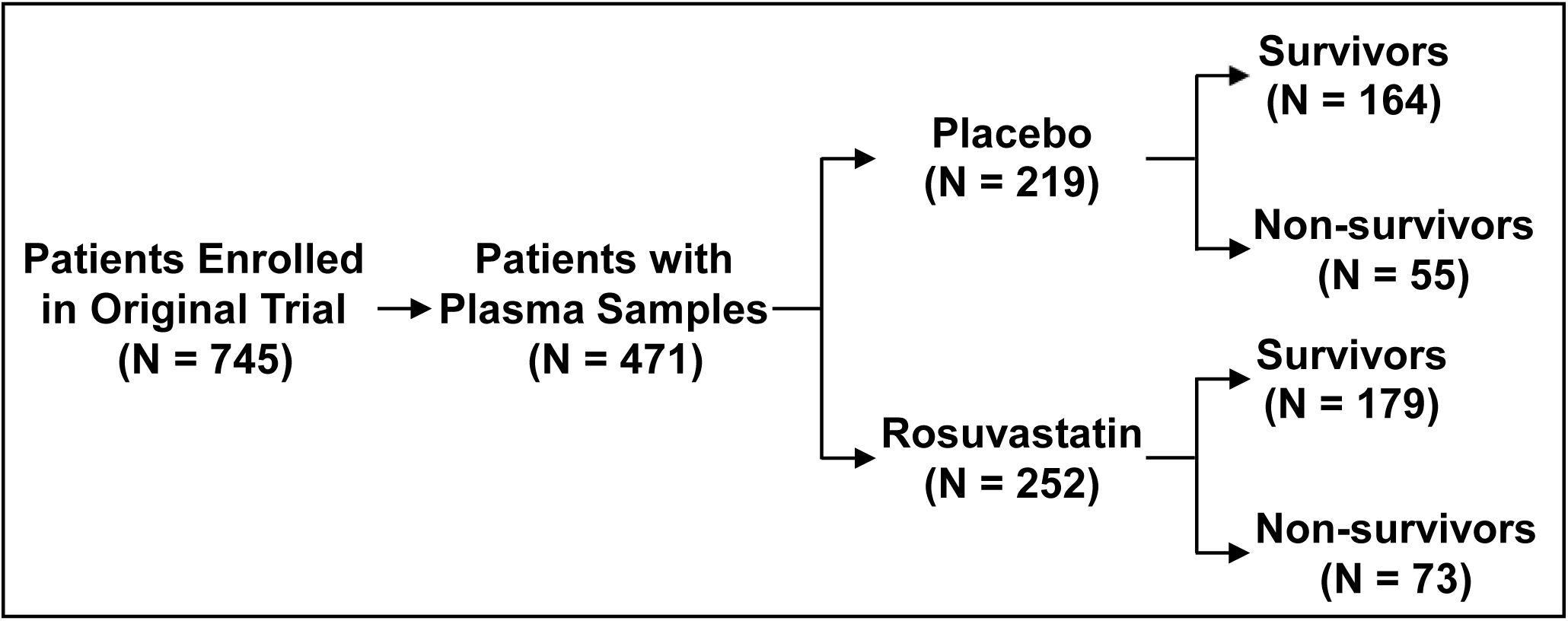
Flow chart of study population in this secondary analysis.

### Mice study

To directly evaluate this protective effect in vivo, we overexpressed human CETP in C57BL/6 mice (these mice do not naturally express CETP) (17) via liver-directed adeno-associated virus (serotype 8) expressing human CETP (AAV8-TBG-h-CETP) (AAV-CETP) delivery. Empty AAV vector (AAV-null) was used as the control. Two weeks post-injection, AAV-CETP mice (5×10^10^ cg/mouse) exhibited elevated plasma CETP activity compared to AAV-null controls (Supplemental Fig. S1A). Upon challenge with lipopolysaccharide (LPS, 7 mg/kg, i.p.) to induce endotoxemia, survival was monitored over 7 days. Mice were housed in micro-isolator cages and maintained on a 14-hr light/10-hr dark cycle. Mice were provided with normal chow diet and water *ad libitum*. All studies were performed in accordance with the Public Health Service Policy on Humane Care and Use of Laboratory Animals and with the approval of the University of Kentucky and Lexington Veterans Affairs Medical Center Institutional Animal Care and Use Committees.

### Plasma Analysis

Plasma CETP activity assay (MAK106, Sigma), and SAA levels (EK249341, AFG Bioscience, Arlington Heights, IL) were assessed using commercially available kits, per the manufacturer’s instructions.

### Fast Protein Liquid Chromatography (FPLC)

Pooled plasma samples were generated from six patients per pool within each study subgroup, including survivors and non-survivors from both the PL and RS treatment arms. A total of six independent pools were analyzed per subgroup (n = 6 pools/group). Plasma pools were fractionated by FPLC using an AKTA Pure instrument equipped with a Superose 6 Increase column (Cytiva) at a flow rate of 0.75 mL/min. Total cholesterol (Pointe Scientific, Canton, MI) concentrations and SAA levels in the collected fractions were quantified using enzymatic assays and ELISA respectively.

### Statistical Analysis

Differences in biomarkers between the PL and the RS groups were assessed over time using linear mixed models to account for repeated measurements within patients. Additionally, survivors and non-survivors in each group were separated for further exploration, and linear mixed models were used to assess temporal within-group differences and timepoint-specific cross-group differences between groups and between survivors and non-survivors. Likelihood ratio testing and Akaike Information Criterion (AIC) were used to select an appropriate covariance structure for each mixed model, and a Kenward-Roger adjustment was used to correct for negative bias in the standard error and degrees of freedom estimation. Log or square-root transformation of the biomarkers was used to correct for right skewness, as appropriate. For all other outcomes that did not involve repeated measurements, separate models were fitted within the PL and RS groups. Cox proportional hazards models were used to assess the relationship between biomarker levels and survival. Linear regression models were used to evaluate the correlation between biomarker levels and APACHE score, with confidence intervals for the correlations calculated using Fisher’s *z*-transformation. Zero-inflated negative binomial models were used to examine the association between biomarker levels and ICU-free days, while beta-binomial regression models were used for bounded count outcomes representing dysfunction-free days.

Pairwise associations among the measured biomarkers were assessed using Spearman correlation analyses due to heavy right skewness in all observed biomarker distributions, with bootstrap 95% confidence intervals included in the reporting.

## Results

This study analyzed clinical data and plasma samples from patients enrolled in the ARDSNet SAILS trial with available outcome data and archived plasma specimens obtained at the day of enrollment (day 0) and study days 3 and 6 from the BioLINCC biorepository. The distribution of available samples is shown in (Fig. 1). In total, 1,233 plasma samples from 471 patients were analyzed for CETP activity and SAA levels. Because not all participants had plasma available at every study time point, the number of samples analyzed varied across days 0, 3, and 6.

### Temporal changes in CETP activities and their association with sepsis survival

There were no significant changes in CETP over time across all groups, except in the PL survivors, where activities were modestly but significantly increased from Day 0 to Day 3 and Day 0 to Day 6 (Mean ± SEM for Days 0, 3 and 6: 6.52±0.46, 7.43±0.47, and 8.02±0.61 respectively). RS treatment did not significantly change CETP activities in both sepsis survivors and non-survivors (Fig. 2A). CETP activities were significantly higher in the survivors compared to non-survivors on all three days of analysis, in both the treatment groups (PL group: p=0.002, p=0.0008 and p=0.010 respectively for Days 0, 3 and 6; for RS group: p=0.006, p=0.003, and p=0.015 for Days 0, 3 and 6 respectively). RS treatment did not change the CETP activities in both survivors and non-survivors (Fig. 2A).

**Fig. 2.**
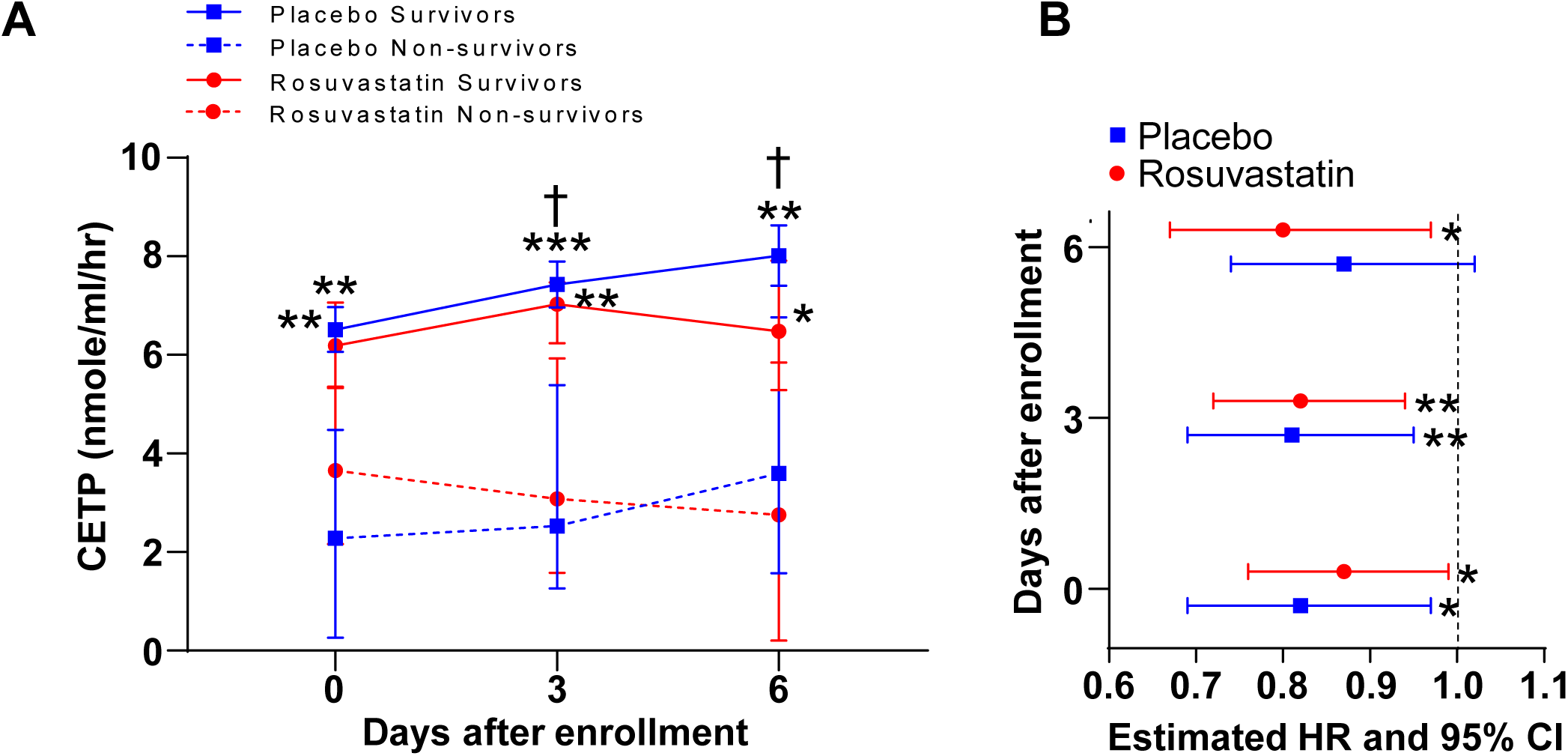
Temporal changes in CETP activities and their association with sepsis survival. **A,** Plasma CETP activities were determined on days 0, 3, and 6 after enrollment, in patients assigned to the placebo and rosuvastatin treatment groups. Means with SEM are shown in the figures. CETP activities were assessed using commercially available kit. Linear mixed models assessed differences in plasma CETP activities over time between survivors and non-survivors from each group. *=p<0.05 versus rosuvastatin non-survivors on the same day; **=p<0.01 versus non-survivors on the same day and same treatment; ***=p<0.001 versus placebo non-survivors on the same day. †=p<0.05 for CETP activities between Day 0 and Day 3 and Day 0 and Day 6. **B,** Associations between plasma CETP activities and death. Plasma CETP activities were determined on days 0, 3, and 6 of enrollment in patients assigned to the placebo and rosuvastatin treatment groups. Separate Cox regression models were fit for each group to assess the association between survival time and CETP activities. Corresponding hazard ratios (HR) and 95% confidence intervals (CI) were shown for each group on each day. Statistical significance is denoted as *=p<0.05, and **=p<0.01.

All odds ratios and hazard ratios from survival analysis (Cox regression) expressed per3-unit increase of CETP activities indicated the following results. In PL patients, an increase in CETP activity on the day of enrollment (Day 0) and Day 3 was positively associated with survival (Day 0: HR 0.82, p=0.021; Day 3: HR 0.81, p=0.008). Likewise, in RS patients, an increase in CETP levels on all three days, Days 0, 3, and 6, was significantly associated with better survival (Day 0: HR 0.87, p=0.039; Day 3: HR 0.82, p=0.005, and Day 6: HR 0.80, p=0.022; Fig. 2B).

Spearman correlation analysis (with 95% bootstrap confidence intervals) of CETP activities with various cholesterol components in plasma, total cholesterol (TC), free cholesterol (FC), and high-density cholesterol (HDL-C) indicated a statistically significant but weak negative correlation (ρ=-0.152, p<0.001) with HDL-C, no significant correlation with plasma total cholesterol levels, and a statistically significant but weak positive correlation with FC, backed by a 95% confidence interval (0.042 to 0.157) (Supplemental Table S1).

### Higher CETP activities associate with better sepsis outcomes in patients treated with RS

Plasma CETP activities had no association with disease severity, including determinants such as APACHE scores or hepatic-, renal-, or overall organ dysfunction-free days in patients in the PL group (Fig. 3). However, in the RS group, higher plasma CETP on days 3 and 6 negatively correlated with APACHE scores (Day 3: r=-0.14, p=0.023 and Day 6: r= −0.25, p<0.001; Fig. 3A). Consistently, patients in the RS group who had higher plasma CETP activities on Days 3 and 6, tended to have more renal dysfunction-free days with the association becoming stronger by Day 6 (Day 3: r=1.13 (95% CI 1.02–1.26), p = 0.023 and Day 6: r=1.29 (95% CI 1.11–1.50), p=0.001; Fig. 3B). However, higher CETP activities on Day 0 was associated with fewer hepatic dysfunction-free days, suggesting worse liver outcomes early on (Day 0: 0.86 (95% CI 0.77– 0.96), p= 0.008; Fig. 3C). Neither treatment group showed statistically significant association with overall organ dysfunction at any time point (Fig. 3D).

**Fig. 3.**
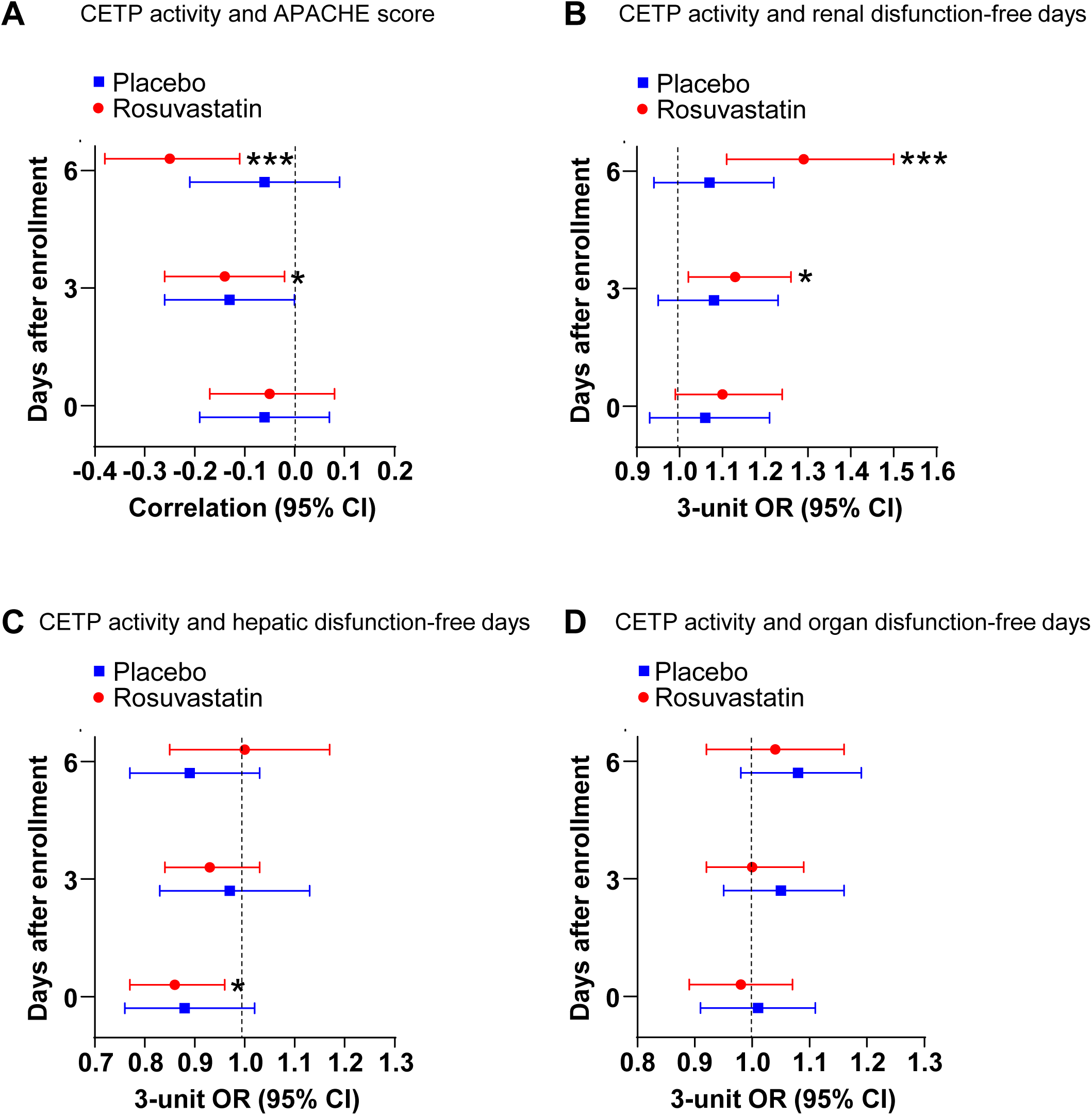
Higher CETP activities associate with better sepsis outcomes in patients treated with rosuvastain. **A,** Associations between plasma CETP activities and APACHE score. Plasma CETP activities were determined on days 0, 3, and 6 of enrollment in patients assigned to the placebo and rosuvastatin treatment groups. Linear regression models were used to evaluate the correlation between CETP activities and APACHE score, with 95% confidence intervals (CI) for the correlations for each group on each day. Statistical significance is denoted as *=p<0.05, and ***=p<0.001. **B,** Plasma CETP activities and renal disfunction-free days analysis. Plasma CETP activities were determined on days 0, 3, and 6 of enrollment in patients assigned to the placebo and rosuvastatin treatment groups. Odds ratios (OR) are expressed per 3-unit (nmole/ml/hr) increase of CETP for interpretability. Statistical significance is denoted as *=p<0.05, and ***=p<0.001. **C,** Plasma CETP activities and hepatic disfunction-free days analysis. Plasma CETP activities were determined on days 0, 3, and 6 of enrollment in patients assigned to the placebo and rosuvastatin treatment groups. Odds ratios are expressed per 3-unit (nmole/ml/hr) increase of CETP for interpretability. **D,** Plasma CETP activities and organ disfunction-free days analysis. Plasma CETP activities were determined on days 0, 3, and 6 of enrollment in patients assigned to the placebo and rosuvastatin treatment groups. Odds ratios are expressed per 3-unit (nmole/ml/hr) increase of CETP for interpretability. For **B/C/D**, the p-values and corresponding ORs and CIs come from separate beta-binomial models for placebo and rosuvastatin group.

### CETP expression improves survival in C57BL/6 mice with endotoxemia

The role of CETP in sepsis remains highly controversial. Clinical studies are conflicting, while some investigators report a positive association between CETP levels and sepsis survival (18), others indicate that CETP negatively impacts survival (9, 19). Preclinical animal studies show a similar divide, with reports demonstrating both detrimental (20) and protective (11) effects of CETP. In the current study, we observed that human sepsis survivors had significantly higher plasma CETP levels (Fig. 2A) and that elevated CETP activity correlated with a reduced hazard ratio for mortality (Fig. 2B). To directly evaluate this protective effect in vivo, we overexpressed human CETP in C57BL/6 mice via liver-directed adeno-associated virus (AAV) delivery. Two weeks post-injection, AAV-CETP mice (5×10^10^ cg/mouse) exhibited elevated plasma CETP activity compared to AAV-null controls (Supplemental Fig. S1A). Upon challenge with lipopolysaccharide to induce endotoxemia, survival was monitored over 7 days. Consistent with our human cohort observations, AAV-CETP mice exhibited significantly improved survival against lethal endotoxemia compared to control mice (Supplemental Fig. S1B), indicating that CETP is protective under these conditions.

### Temporal changes in plasma SAA levels and their association with sepsis survival

Plasma SAA levels declined substantially from day 0 to days 3 and 6 in both treatment groups (P<0.0001; Fig. 4A). However, in the PL group, survivors had significantly higher Day 0 SAA levels than non-survivors, a difference not observed with RS treatment (Fig. 4B). In the PL group, elevated plasma SAA on Days 0 and 3 was associated with a trend toward increased survival. Conversely, in the RS group, higher SAA levels were associated with an increased hazard ratio, with Day 6 SAA levels demonstrating a significant increase in mortality risk (HR = 1.04; Fig. 4C).

**Fig. 4.**
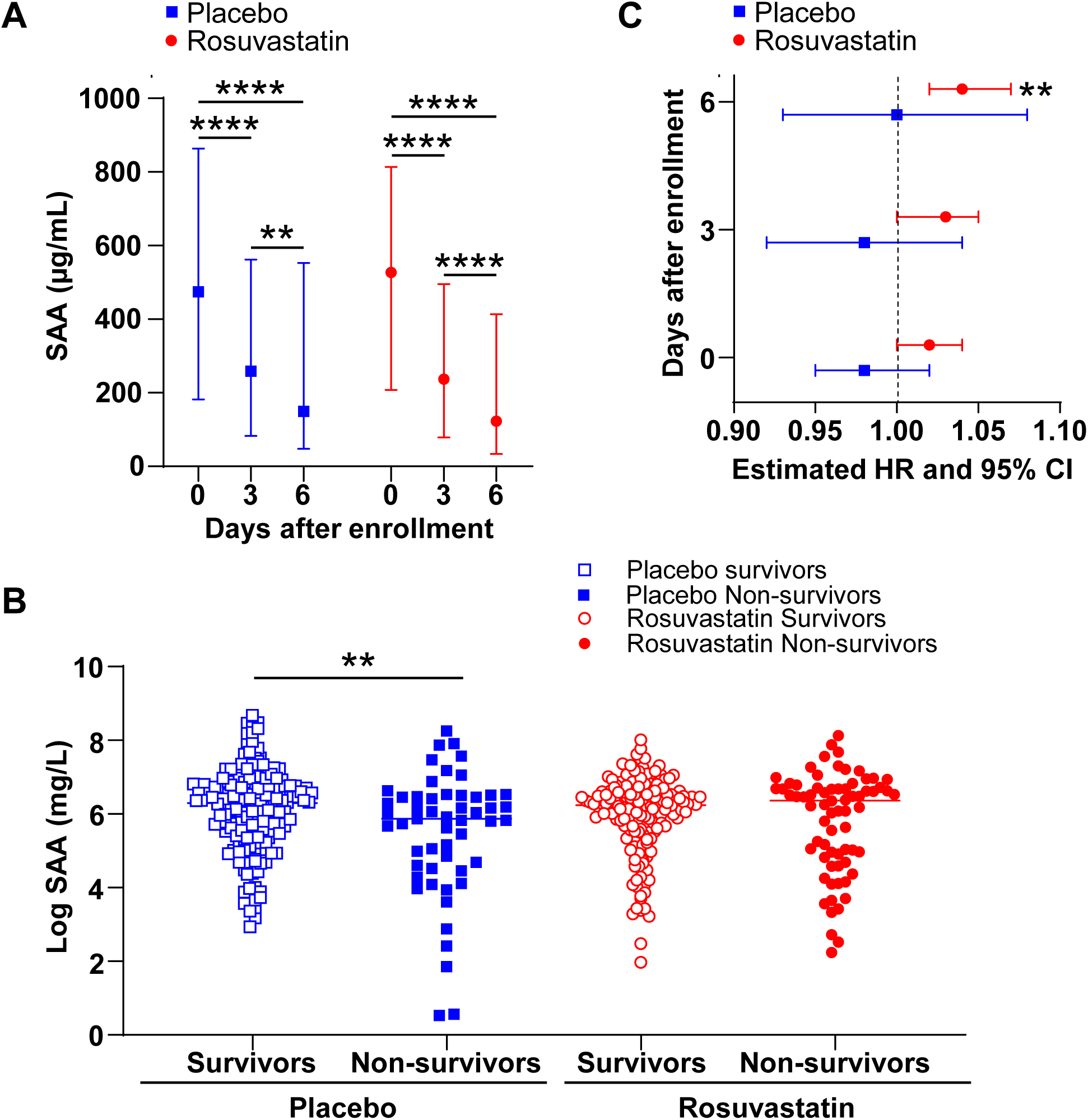
Temporal changes in plasma SAA levels and their association with sepsis survival. **A,** Plasma SAA levels were determined on days 0, 3, and 6 of enrollment in patients assigned to the placebo and rosuvastatin treatment groups. Means with SEM are shown in the figures. SAA levels were assessed by ELISA using commercially available kit. Statistical significance is denoted as **=p<0.01, and ****=p<0.0001. Linear mixed models assessed differences in plasma SAA levels over time. **B,** Plasma levels of SAA (day 0) were measured in survivors and nonsurvivors with sepsis in both placebo and rosuvastatin treatment groups. Pairwise comparisons from one-way ANOVA on log-transformed values were used to assess survivors vs. non-survivors. Statistical significance is denoted as **=p<0.01. **C,** Associations between plasma SAA levels and death. Plasma SAA levels were determined on days 0, 3, and 6 of enrollment in patients assigned to the placebo and rosuvastatin treatment groups. Separate Cox regression models were fit for each group to assess the association between survival time and SAA levels. Corresponding hazard ratios (HR) and 95% confidence intervals (CI) were shown for each group on each day. Statistical significance is denoted as **=p<0.01.

### Associations between plasma SAA and clinical outcomes in sepsis

In the PL group, higher SAA was associated with slightly lower APACHE scores at Day 0 and Day 3, in the RS group, this association was observed only on Day 3 (Fig. 5A). Plasma SAA levels on Days 0,3 and 6 did not show any statistically significant association with renal dysfunction-free days. However, among patients who received RS treatment, higher SAA levels at Days 3 and 6 were associated with better renal outcomes, increased renal dysfunction-free days (Day 3: p=0.004 and Day 6: p=0.006; Fig. 5B). Consistent with status of renal dysfunction-free days, plasma SAA levels for patients in the PL group on Days 0, 3 or 6, did not show any significant association with hepatic dysfunction-free days, while patients in the RS group, higher SAA was consistently associated with more hepatic dysfunction-free days and the strength of this association increased over time (Day 0: p=0.001; Days 3 and 6: p<0.001; Fig. 5C). Higher plasma SAA levels were also associated with an increase in overall organ dysfunction-free days in the RS group on all three days of analysis, with a stronger association over time. In the PL group, this relationship was present only on Day 0 (p=0.002; Fig. 5D). The association of plasma SAA levels with clinical outcomes is stronger in the RS group, where higher SAA levels are associated with more renal, hepatic, and overall organ dysfunction-free days, particularly at Days 3 and 6 (Fig. 5B-D).

**Fig. 5.**
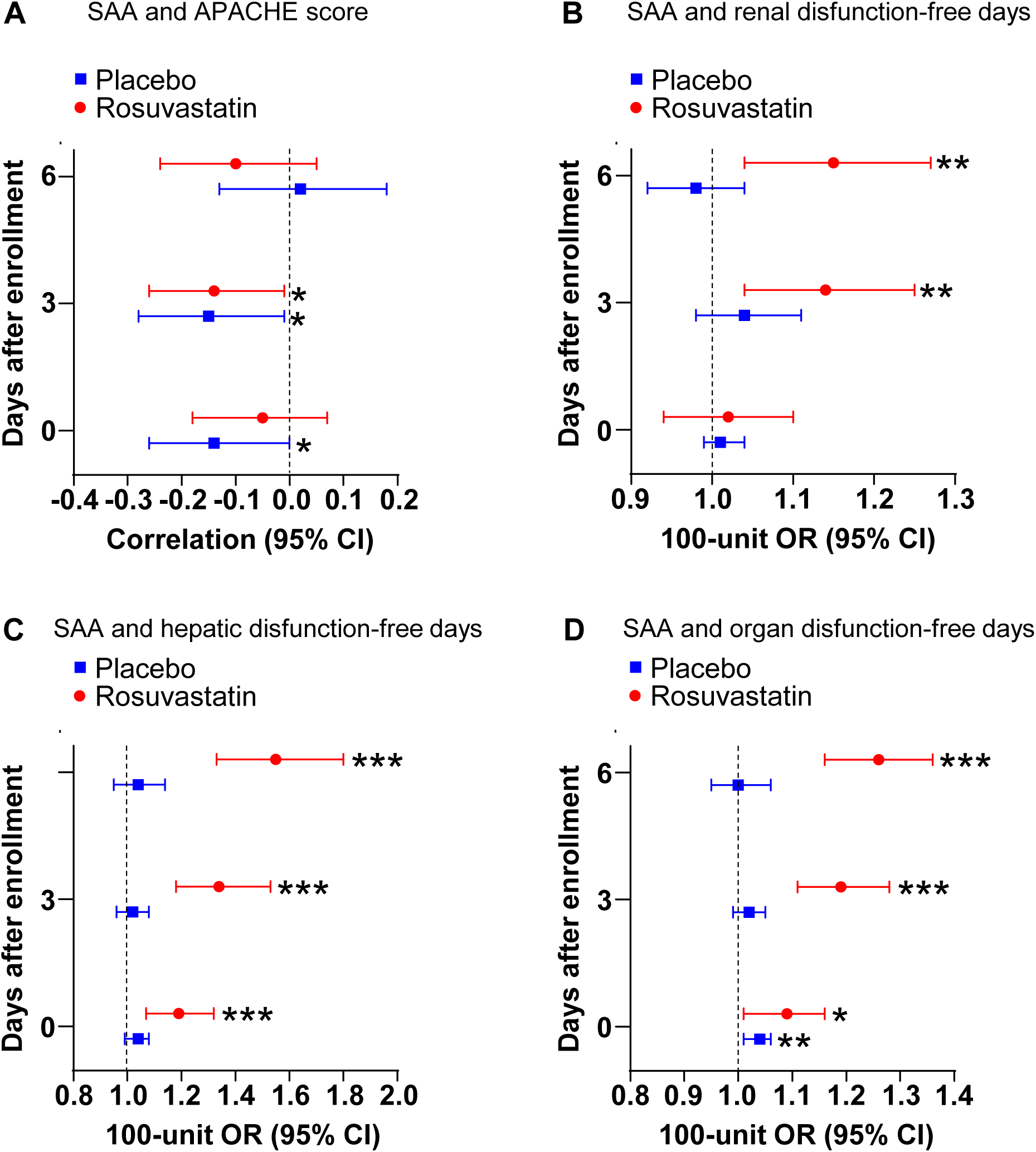
Associations between plasma SAA and clinical outcomes in sepsis. **A,** Associations between plasma SAA levels and APACHE score. Plasma SAA levels were determined on days 0, 3, and 6 of enrollment in patients assigned to the placebo and rosuvastatin treatment groups. Linear regression models were used to evaluate the correlation between SAA levels and APACHE score, with 95% confidence intervals (CI) for the correlations for each group on each day. Statistical significance is denoted as *=p<0.05. **B,** Plasma SAA levels and renal disfunction-free days analysis. Plasma SAA levels were determined on days 0, 3, and 6 of enrollment in patients assigned to the placebo and rosuvastatin treatment groups. Odds ratios (OR) are expressed per 100-unit (µg/mL) increase of SAA for interpretability. Statistical significance is denoted as **=p<0.01. **C,** Plasma SAA levels and hepatic disfunction-free days analysis. Plasma SAA levels were determined on days 0, 3, and 6 of enrollment in patients assigned to the placebo and rosuvastatin treatment groups. Odds ratios are expressed per 100-unit (µg/mL) increase of SAA for interpretability. Statistical significance is denoted as ***=p<0.001. **D,** Plasma SAA levels and organ disfunction-free days analysis. Plasma SAA levels were determined on days 0, 3, and 6 of enrollment in patients assigned to the placebo and rosuvastatin treatment groups. Odds ratios are expressed per 100-unit (µg/mL) increase of SAA for interpretability. Statistical significance is denoted as *=p<0.05, **=p<0.01, and ***=p<0.001. For **B/C/D**, the p-values and corresponding ORs and CIs come from separate beta-binomial models for placebo and rosuvastatin group.

### Survival in sepsis was associated with higher plasma SAA on HDL

Because SAA is predominantly transported on HDL during inflammation, we examined SAA distribution among lipoprotein fractions. Most circulating SAA was recovered in the HDL fraction (Fig. 6A-D). Among PL-treated patients, HDL-associated SAA levels were significantly higher in survivors than in non-survivors on days 0 (p=0.006; Fig. 6A) and 3 (p=0.005; Fig. 6B). This difference was not observed in the RS group (Fig. 6C, D). Among survivors in both treatment groups, SAA appeared to be preferentially associated with denser HDL particles, as indicated by a rightward shift in the distribution curve; this shift was more pronounced in the RS group (Fig. 6A-D).

**Fig. 6.**
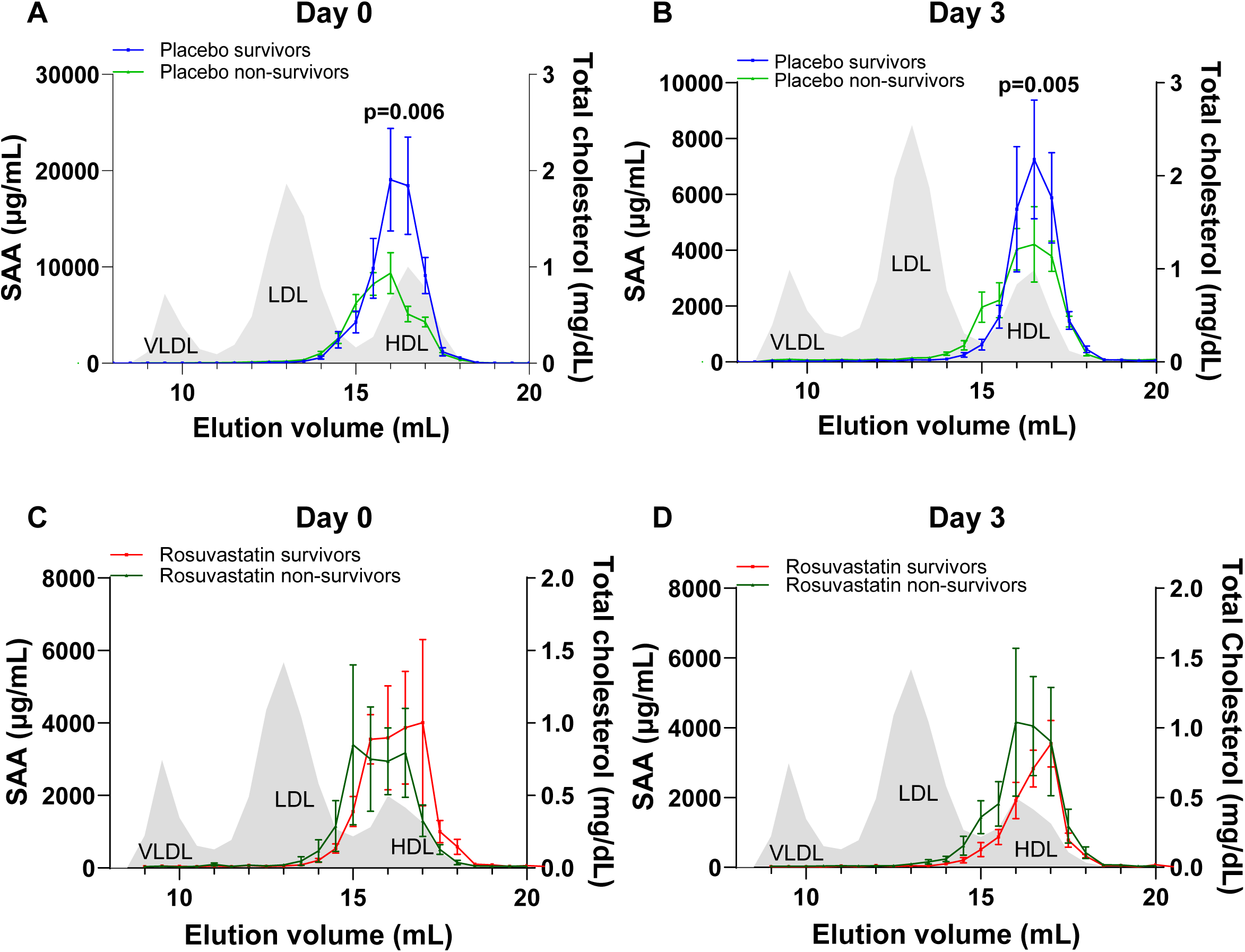
Survival in sepsis was associated with higher plasma SAA on HDL. SAA levels in FPLC-fractionated plasma from sepsis patients in the placebo group of Day 0 (**A**) and Day 3 (**B**) and rosuvastatin group of Day 0 (**C**) and Day 3 (**D**). The indicated p-values were derived from t-tests performed on the area under the HDL peak between survivors and non-survivors. A total of six independent pools were analyzed per subgroup (n=6 pools/group). The mean differences were calculated as non-survivors vs survivors from each group on the same day.

## Discussion

In this study, we performed a longitudinal analysis of CETP activity and SAA, two proteins involved in lipoprotein metabolism and the host response to inflammation (21–23), using plasma samples collected from patients with sepsis-associated ARDS enrolled in the ARDSNet SAILS clinical trial. The SAILS trial was a multicenter randomized controlled trial designed to evaluate the efficacy of RS as a therapeutic intervention for sepsis-associated ARDS. Leveraging these archived longitudinal samples, we investigated temporal changes in CETP activity and SAA levels and their associations with clinical outcomes and RS treatment. RS is a potent HMG-CoA reductase inhibitor that effectively lowers LDL cholesterol (LDL-C) with relatively modest effects on HDL-C. In addition to its lipid-lowering properties, RS exerts pleiotropic effects, including anti-inflammatory, antioxidant, and endothelial-protective actions that are thought to contribute to its cardiovascular benefits beyond LDL-C reduction (24, 25). The study also aimed to evaluate the effect of RS on the lipid-associated proteins CETP and SAA, which influence cholesterol homeostasis in the circulation (21, 23), and to determine whether treatment modifies their association with sepsis outcomes. We show that plasma CETP activity and SAA are dynamically altered during critical illness and are differentially associated with survival, organ dysfunction, and RS exposure. Collectively, these findings support the concept that maintaining homeostasis of these parameters during sepsis may represent an adaptive host response associated with improved recovery.

There was a significant and clear difference in the activities of CETP between survivors and non-survivors of sepsis in both PL and RS groups, with survivors having significantly higher CETP activities compared to non-survivors, especially on Day 3. Consistently, higher CETP activities were significantly associated with improved survival in both the treatment groups, with a stronger association on Day 3. Intriguingly, in the RS group, higher CETP was significantly associated with lower APACHE scores and more renal dysfunction-free days at Days 3 and 6, whereas no statistically significant associations were observed in the PL group.

The role of CETP in sepsis remains incompletely understood, but our findings suggest that preserved CETP activity may be associated with favorable outcomes. In this study, higher CETP activity was consistently associated with survival, particularly in RS-treated patients, and correlated with decreased APACHE scores and increased renal dysfunction-free days. Our findings in patients with sepsis are supported by our study in CETP-overexpressing mice, in which CETP overexpression confers protection against endotoxemia-induced mortality. CETP facilitates the transfer of cholesteryl esters and triglycerides among circulating lipoproteins and plays a central role in lipoprotein remodeling (26). Consistently, we observed a significant negative correlation between HDL-C and CETP activities in this study. Although the role of CETP in sepsis remains controversial, with several studies suggesting detrimental effects (19, 20, 27), others support a protective role for CETP in the septic response (11, 18, 28).

A majority of observational studies reported a negative correlation between HDL concentration and mortality (29, 30). An apparent paradox arises from the observation that both higher HDL-C levels and preserved CETP activity are associated with improved sepsis outcomes, despite CETP activity negatively correlating with HDL-C levels (Supplemental Table S1). Our findings suggest that CETP activity itself may confer beneficial effects in sepsis that are independent of, or may partially offset, its HDL-C–lowering properties. Consequently, therapeutic strategies aimed at inhibiting CETP to raise HDL-C levels, such as repurposing CETP inhibitors that failed to prove effectiveness in reducing cardiovascular disease risk, in sepsis may produce unintended adverse effects. Taken together, these findings indicate that maintaining a balance in CETP activity may be critical, preserving sufficient CETP function to retain its protective effects while avoiding excessive reductions in HDL-C levels that could compromise host defense and survival in sepsis. It is important to note that the CETP inhibitor (torcetrapib) significantly increased death from infections in the Atherosclerotic Events trial, ILLUMINATE (31), however, other trials, dalcetrapib (32), anacetrapib (33) or evacetrapib (34) did not report this observation.

Beyond their lipid-lowering effect, statins also have pleiotropic properties including anti-inflammatory, antioxidant, immunomodulatory, and antithrombotic effects (35–37), these properties were believed to offer sepsis protection. The original study reported a significant but weak positive correlation between the CRP level with the RS treatment (4). SAA, like CRP is an acute-phase protein, in this study, we did not observe any significant effects of RS on SAA levels. SAA levels reduced significantly with time in both the treatment groups. Using preclinical mouse models, our group has reported that SAA is protective in sepsis, with SAA deficiency significantly increasing sepsis mortality (16). Interestingly, we observed that higher SAA levels on Day 0 were associated with survival in PL-treated patients. Consistent with its association with survival, early increases in plasma SAA levels also negatively correlated with APACHE score, an indicator of sepsis severity. The original ARDs SAILs study reported RS therapy to be associated with fewer days free of renal and hepatic failure to day 14 (4). In this study we observed that patients in the RS treatment groups with higher plasma SAA have significantly higher number of days free of renal and hepatic dysfunction. All these results indicate that early increases in SAA levels possibly plays a protective role in sepsis and anti-inflammatory therapy which decreases plasma SAA levels may negatively affect the prognosis of the disease.

SAA as a potential mediator of HDL’s protective effects in sepsis: During inflammation, SAA becomes one of the major apolipoproteins associated with HDL (23), with virtually every HDL particle reported to contain at least one SAA molecule (38). Thus, HDL appears to serve as an important circulating carrier for SAA, facilitating its mobilization and accumulation during the acute-phase response. Notably, low circulating HDL levels have been associated with increased risk and severity of sepsis, and strategies aimed at increasing HDL levels have been proposed as potential therapeutic approaches (39). Given that apolipoproteins are major structural and functional components of HDL, these observations raise the important question of whether SAA contributes to the biological properties of HDL during sepsis. In the present study, we found that the majority of circulating SAA was associated with the HDL fraction and that SAA levels were significantly higher among survivors than non-survivors in the PL group. Although these findings do not establish a causal relationship, they support the hypothesis that SAA may contribute to the protective functions of HDL during sepsis. Accordingly, a reduction in circulating SAA-HDL may reflect impaired HDL-associated protective mechanisms and could be associated with worse clinical outcomes in sepsis.

Our study has several important strengths, including the large number of longitudinal plasma samples and the use of a well-characterized, multicenter ARDS cohort assessed at several clinically relevant time points. The integration of SAA concentrations and CETP activity measurements with survival and organ dysfunction outcomes provides a comprehensive characterization of the temporal dynamics of SAA and CETP in sepsis-associated ARDS. Several limitations should also be acknowledged. First, this was a retrospective biomarker analysis of a completed clinical trial; therefore, causal relationships cannot be established. Second, plasma sample availability varied across time points, potentially introducing selection bias. Third, the study primarily evaluated circulating SAA concentrations and CETP activity and did not directly assess their biological functions. In addition, residual confounding related to illness severity, nutritional status, liver function, and concomitant therapies may have influenced the observed associations. Finally, because the study population consisted of patients with sepsis-associated ARDS, the findings may not be generalizable to broader populations of patients with sepsis.

In conclusion, this study demonstrates that preservation of plasma CETP activity and higher early SAA levels during sepsis-associated ARDS are associated with improved survival and reduced organ dysfunction. Although RS treatment did not significantly affect CETP activity or SAA levels, it appeared to modify the association between SAA and mortality. These findings support further investigation into the biological roles of SAA and CETP in critical illness and their potential contributions to outcomes in sepsis.

## Authors’ contributions

MES and PS: conceptualization; AJ, OA, LWM, HY, CHV, MECB and SMG: investigation and plasma lipids analysis; GSH: data analysis and interpretation; AJ and PS: original draft; AJ, MES and PS: review and editing; MES and PS: funding acquisition.

## Ethics approval and consent to participate

The original SAILS clinical trial (4) (ClinicalTrials.gov Identifier: NCT00979121, accepted September 16, 2009) was approved by the Institutional Review Board at each of the 44 enrolling hospitals in the NHLBI ARDS Clinical trial network, and assent for participation was collected from patients and their families as part of SAILS trial enrollment at all centers. All the patients or their representatives provided written informed consent or assent at all centers of the trial, including consent to use biospecimens for future research for subjects included in this analysis, and the study was conducted in accordance with the principles of the Declaration of Helsinki. The current study was reviewed by the Institutional Review Board at the University of Kentucky and received a Not Human Research (NHR) determination; therefore, IRB approval was not required. The NHR determination was submitted to the NIH BioLINCC prior to the release of the data and samples.

## Human Ethics and Consent to Participate declarations

Not applicable.

## Availability of data and materials

The datasets used and/or analyzed during the current study are available from the corresponding author on reasonable request.

