## Supplemental Materials for "Preserved CETP Activity and Early Serum Amyloid A are Associated with Improved Outcomes in Sepsis-Associated ARDS"

| Plasma activity | Plasma Levels | Spearman ρ (Bootstrap 95% CI) | P-value |
|---|---|---|---|
| CETP | HDL-C | -0.152 (-0.207 to -0.096) | < 0.001 |
| CETP | TC | 0.048 (-0.009 to 0.105) | 0.094 |
| CETP | FC | 0.100 (0.042 to 0.157) | < 0.001 |

**Supplemental Table S1.** Spearman correlation analysis with 95% bootstrap confidence intervals revealed a statistically significant but weak negative correlation between CETP activity and HDL-C levels ($\rho = -0.152$, $p < 0.001$). CETP activity was not significantly correlated with plasma total cholesterol (TC) levels but showed a statistically significant, weak positive correlation with free cholesterol (FC), with a 95% bootstrap confidence interval of 0.042 to 0.157.

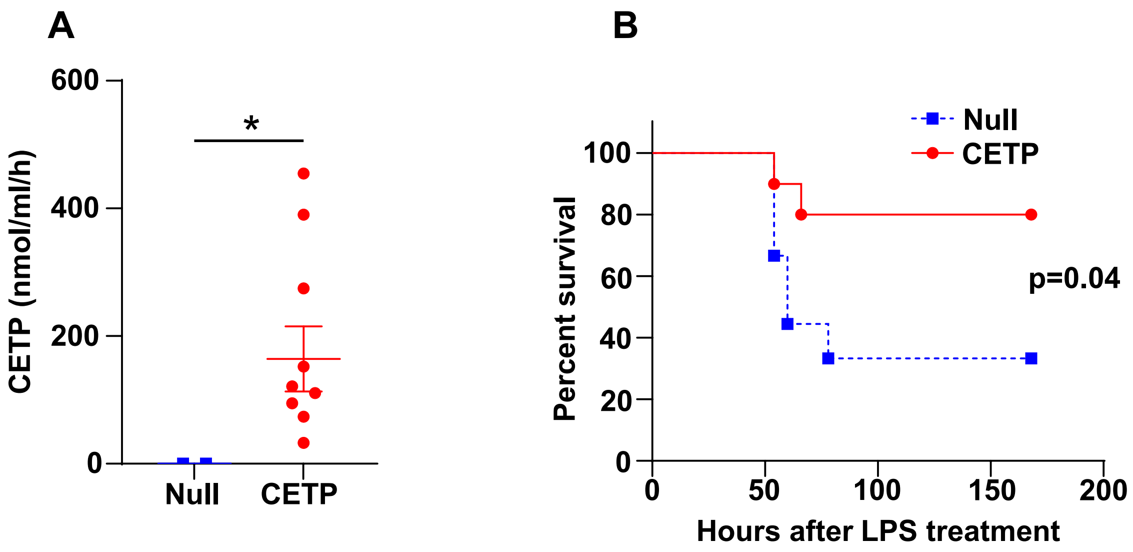


**Supplemental Fig. S1. CETP expression improves survival in C57BL/6 mice with endotoxemia. A,** CETP was expressed in C57BL/6 mice, which do not naturally express CETP, via liver-directed delivery of an adeno-associated virus serotype 8 vector expressing human CETP (AAV8-TBG-h-CETP; $5x10^{10}$ cg/mouse). Mice receiving an empty AAV vector (Null) served as controls. Two weeks after injection, plasma CETP activity was assessed using a commercially available assay kit (MAK106, Sigma-Aldrich) according to the manufacturer's instructions. **B,** CETP male mice (n = 10) and Null control male mice (n = 9) were challenged with lipopolysaccharide (LPS; 7 mg/kg, i.p.) to induce endotoxemia. Survival was monitored for 7 days following LPS administration. Mice were housed in micro-isolator cages and maintained on a 14-hr light/10-hr dark cycle. Mice were provided with normal chow diet and water *ad libitum*. All studies were performed in accordance with the Public Health Service Policy on Humane Care and Use of Laboratory Animals and with the approval of the University of Kentucky and Lexington Veterans Affairs Medical Center Institutional Animal Care and Use Committees.
